# Forward and Reverse Genetic Dissection of Morphogenesis Identifies Filament-Competent *Candida auris* Strains

**DOI:** 10.1101/2021.04.29.442079

**Authors:** Darian J. Santana, Teresa R. O’Meara

## Abstract

*Candida auris* is an emerging healthcare-associated pathogen of global concern. Although this organism does not display the same morphological plasticity as the related fungal pathogen *Candida albicans*, recent reports have identified numerous *C. auris* isolates that grow in cellular aggregates or filaments. However, the genetic circuitry governing *C. auris* morphology remains largely uncharacterized. Here, we developed an *Agrobacterium-mediated* transformation system to generate mutants exhibiting aggregating or filamentous cell morphologies. Aggregating strains were associated with disruption of homologs of *Saccharomyces cerevisiae* chitinase and chitin synthase regulatory proteins, including components of the Regulation of *ACE2* Morphogenesis (RAM) pathway, while disruption of a homolog of the *S. cerevisiae ELM1* gene resulted in a novel filamentous strain of *C. auris*. To facilitate targeted genetic manipulation, we developed a transiently expressed Cas9 and sgRNA expression system for use in *C. auris*. Transformation using this system significantly increased the efficiency of homologous recombination and targeted integration of a reporter cassette in all four clades of *C. auris*. Using this system, we generated targeted deletion mutants to confirm the roles of RAM and Elm1 proteins in regulating *C. auris* morphogenesis. Overall, our findings provide novel insights into the genetic regulation of aggregating and filamentous morphogenesis in *C. auris*. Furthermore, the genetic manipulation tools described here will allow for inexpensive and efficient manipulation of the *C. auris* genome.

**Importance:** *Candida auris* is an emerging and often multi-drug resistant fungal pathogen responsible for outbreaks globally. Current difficulties in performing genetic manipulation in this organism remain a barrier to understanding *C. auris* biology. Homologous recombination approaches can result in less than 1% targeted integration of a reporter cassette, emphasizing the need for new genetic tools specific for manipulating *C. auris*. Here, we adapted Agrobacterium-mediated transformation and a transient Cas9 and sgRNA expression system for use in forward and reverse genetic manipulation of *C. auris*. We demonstrated the efficacy of each system by uncovering genes underlying cellular morphogenesis in *C. auris*. We identified a novel filamentous mutant of *C. auris*, demonstrating that this organism has maintained the capacity for filamentous growth. Our findings provide additional options for improving the genetic tractability of *C. auris*, which will allow for further characterization of this emerging pathogen.

## Introduction

Since its 2009 isolation from the ear canal of a patient in Japan, the emerging fungal pathogen *Candida auris* has caused infections and outbreaks in at least 44 countries on 6 continents (2). The global prevalence of *C. auris* is characterized by the seemingly simultaneous emergence of four distinct genetic clades, differing on the scale of hundreds of thousands of single nucleotide polymorphisms (SNPs), with a potential fifth clade recently identified (3, 4). Individual isolates exhibit significant heterogeneity both within and between clades in murine models of infection and colonization (5, 6). The continually increasing understanding of biologically and clinically relevant phenotypic variation among *C. auris* isolates, and the variation between *C. auris* and other well-studied model organisms, emphasizes the need for facile genetic manipulation approaches to allow for mechanistic characterization of this organism.

Although *C. auris* does not form filaments under many of the same environmental cues that induce hyphal growth in *C. albicans* (7), numerous reports of irregular or multicellular growth indicate *C. auris* does exhibit cellular polymorphism. Depletion of the essential molecular chaperone *HSP90* and genotoxic stress induced by hydroxyurea result in elongated cell growth (7, 8). Growth in high salt concentrations induces cell elongation (9). Strains exhibiting filamentous, elongated, or aggregating morphologies have also been isolated from populations of *C. auris* cells following murine infection (10, 11). Numerous reports detail patient isolates with multicellular aggregating properties, often described by a failure of cell aggregates to disperse upon mixing or vortexing (12–15). Aggregating isolates exhibit reduced biomass in biofilm formation and lower virulence in *Galleria mellonella* infection models compared to non-aggregating counterparts (12, 16). Still, the genetic determinants of irregular morphogenesis in *C. auris* remain largely unexplored due in part to difficulties in performing genetic manipulation in this organism.

Transformation of *C. auris* is complicated by low rates of targeted integration and variable transformation efficiency among isolates and clades. The use of RNA-protein complexes of purified Cas9 and gene-specific guide RNAs, referred to as Cas9-ribonucleoproteins (RNPs), to promote homology directed repair demonstrably increases transformation efficiency and targeted integration rates (17). Transformation incorporating RNPs is often the method of choice for manipulating the *C. auris* genome, and variations exist using multiple gRNA target sites to further improve targeted integration efficiency (18). The use of RNPs in transformation, however, comes with increased expense and additional technical considerations during transformation. In *Candida albicans*, transformation with linearized gene cassettes encoding Cas9 and sgRNA promote homozygous gene deletion; these cassettes cannot be detected in the genome of transformants, suggesting they are transiently expressed and not stably integrated (19). A similar transiently expressed CRISPR-Cas9 system promotes targeted genetic manipulation in *Cryptococcus neoformans* (20). We hypothesized that specific adaptation of the transiently expressed CRISPR-Cas9 system to use *C. auris*-recognized promoters would increase the rates of targeted transformation efficiency.

A forward genetics system represents an alternative approach for manipulating the genome. *Agrobacterium tumefaciens-mediated* transformation (AtMT) is an insertional mutagenesis approach with a history of proven success in fungal species (21). *A. tumefaciens* is a plant pathogen that causes crown gall in dicotyledonous plants through genetic transformation (22). Its capacity for transformation is not limited to plants and can be taken advantage of to perform insertional transformation in a variety of eukaryotic species, including *C. albicans, Candida glabrata*, and *Saccharomyces cerevisiae* (23). In practice, mobilization of a DNA sequence flanked by left and right direct repeats (T-DNA) is accomplished by induction of virulence genes during co-culture with a recipient organism using acetosyringone (24). This T-DNA sequence is encoded on the Ti Plasmid harbored by *A. tumefaciens* and can be manipulated to contain fungal selectable markers.

We used AtMT to generate an insertional mutant library in *C. auris* and identified morphogenic mutants growing in aggregates or as pseudohyphae. Insertions in genes orthologous to regulators of chitinase and chitin synthase in *S. cerevisiae* were associated with defects in daughter cell separation in *C. auris*, leading to aggregating growth, while an insertion in an ortholog of *ScELM1* resulted in constitutive filamentous growth in *C. auris*. We developed a robust transient CRISPR-Cas9 expression system for *C. auris* and demonstrated its ability to significantly increase targeted transformation in isolates from all four major clades. Using this system, we performed deletions in key regulators of cell separation to demonstrate functional conservation of *ELM1* and regulators of *ACE2* in *C. auris*. The tools presented here allow for novel analyses of the genetic circuitry required for morphogenesis in the emerging pathogen *C. auris* and will serve as a resource to the community for future molecular genetic manipulation of this pathogen.

## Results

### *Agrobacterium-mediated* transformation identifies *C. auris* morphogenic mutants

While aggregating and filamentous strains of *C. auris* have been recovered from human and murine hosts, the genetic circuitry governing *C. auris* morphogenesis remains largely uncharacterized. Therefore, we set out to apply a forward genetic approach to identify regulators of morphogenesis in *C. auris*. To accomplish this, we developed an *Agrobacterium tumefaciens-mediated* transformation (AtMT) system for *C. auris*. We cloned the CaNAT1 nourseothricin resistance cassette into the pPZP Ti plasmid backbone between the T-DNA left and right borders to generate pTO128 (pPZP-NATca) and transformed the resulting vector into *A. tumefaciens* strain EHA105, which also harbors the virulence genes necessary for mobilization of the T-DNA. We chose the South Asian (Clade I) *C. auris* isolate AR382 from the FDA-CDC Antimicrobial Resistance Isolate Bank as the genetic background for insertional mutagenesis (25). We co-cultured *A. tumefaciens* and *C. auris* on Induction Media containing acetosyringone to induce mobilization of T-DNA and identified transgene insertional mutants of *C. auris* by selecting on YPD plates containing nourseothricin. Because previous reports suggest that AtMT transformation efficiency in fungi varies with alterations in co-culture and incubation parameters (26), we monitored the transformation efficiency of *C. auris* after 2, 4, and 7 days of co-culture incubation with different ratios of *C. auris* and *A. tumefaciens* inocula (Fig. 1A). By comparing the rate of transformants to input, we calculated a maximum transformation efficiency of approximately 1 in 6500 *C. auris* cells at 4 days of co-incubation with equal inocula, which is consistent with the range of transformation efficiencies exhibited in integrative AtMT of other yeast species (26, 27). We performed AtMT in *C. auris* AR382 using these optimal co-culture parameters and identified 6 mutants with altered colony morphology, suggestive of an alteration in cellular morphology (Fig. 1B). These findings demonstrate the utility of AtMT as a forward genetics system for *C. auris*.

**Fig. 1.**
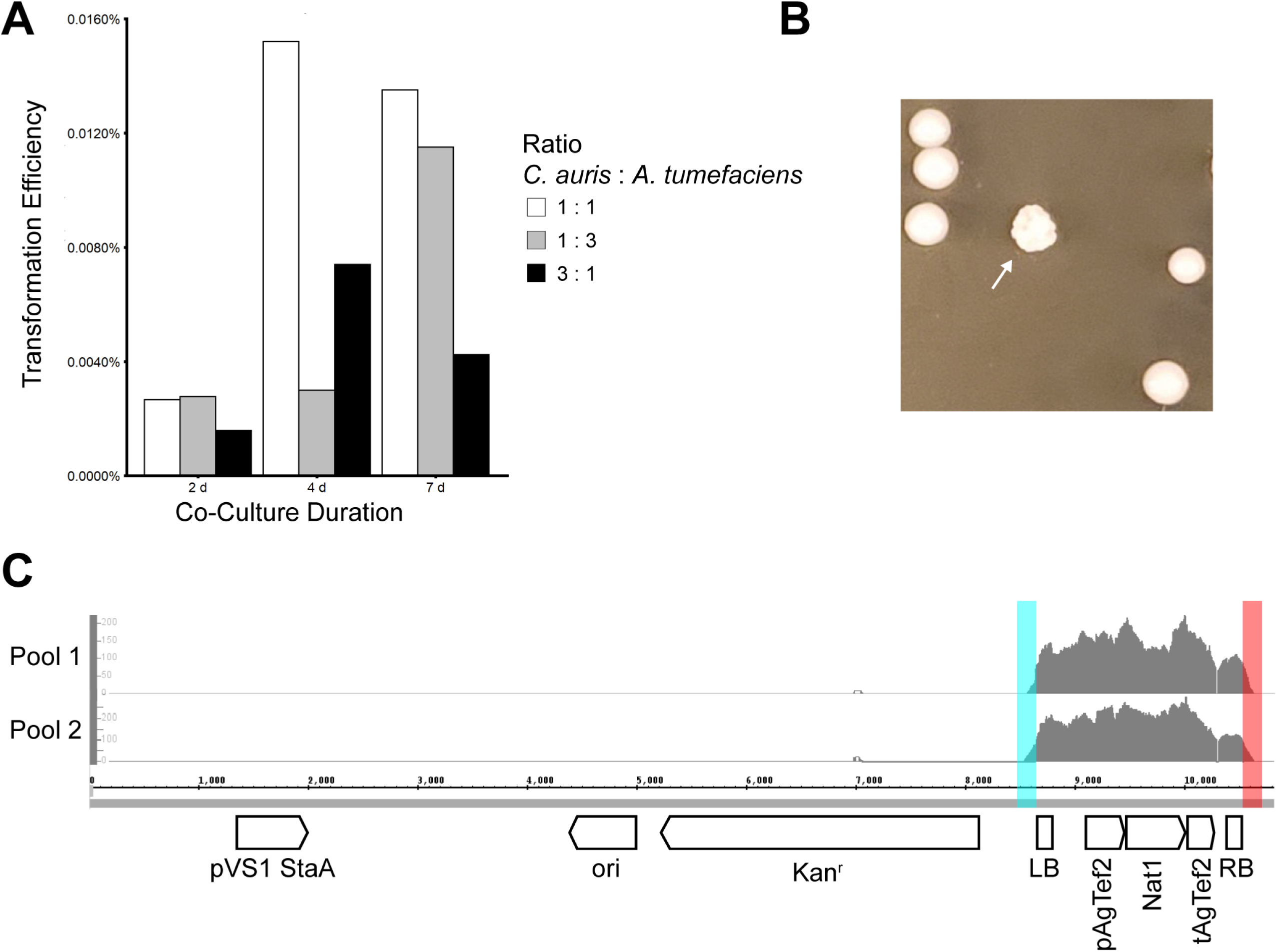
*Agrobacterium tumefaciens-mediated* transformation (AtMT) identifies regulators of colony morphology in *C. auris*. (A) AtMT transformation efficiency of *C. auris* was measured after 2, 4, and 7 days of coculture with three different combinations of *C. auris* to *A. tumefaciens* inocula. Transformation efficiency is expressed as the ratio of recovered *C. auris* transformants to the total number of input *C. auris* cells. (B) Morphogenic mutants were identified in *C. auris* AtMT transformants through irregular colony morphologies (arrow). (C) Genomic DNA was extracted from 6 morphogenic mutants and pooled into two pools of 3 for Illumina sequencing. Reads were mapped to the TI Plasmid (pTO128). Highlighted regions in blue and red indicate read sequence that extended beyond the T-DNA left and right borders, respectively, used to identify transgene insertion sites in the *C. auris* genome.

Transgene insertion sites can be defined by identifying the genomic regions flanking the insertions using whole-genome sequencing (28). We reasoned a similar approach could identify transgene insertion sites from multiple mutants sequenced in pools. We generated 150 bp paired-end Illumina sequencing reads from two pools of three morphogenic mutants each and mapped the sequencing reads from each pool to the sequence of the TI plasmid pTO128 (pPZP-NATca). The sequencing reads mapped exclusively to the T-DNA region of the plasmid, with additional read length spanning either junction at the T-DNA left and right borders (Fig. 1C). The sequence extending beyond the left and right borders corresponded to *C. auris* genomic regions flanking the transgene insertions. We extracted the sequence data from these regions and generated consensus sequences based on multiple sequence alignments. We then mapped the consensus sequences to the *C. auris* B8441 reference genome (NCBI GCA_002759435.2, South Asian Clade) to identify insertion sites and determined which morphogenic mutant harbored which specific insertion using insertion site-specific PCR and Sanger sequencing.

Among the mutants identified by irregular colony morphologies, four exhibited a similar aggregating phenotype, with individual cells connected into clusters that could not be disrupted by vortexing (Fig. 2). We also observed several cell compartments containing more than one nucleus, suggesting a defect in nuclear separation associated with this failure of cell separation. Insertion events in *CauACE2* (B9J08_000468), orthologous to *S. cerevisiae ACE2* (YLR131C), as well as in *CauTAO3* (B9J08_000181), orthologous to *S. cerevisiae TAO3* (YIL129C), were associated with this aggregatory phenotype. A similar aggregating phenotype resulted from an insertion near the C-terminus of *CauCHS2* (B9J08_003879), an ortholog of *CHS2* (YBR038W) in *S. cerevisiae*. A fourth aggregating strain was associated with an insertion in the promoter region of *B9J08_002252;* however, orthologs of this gene in related species are poorly characterized. To predict a potential function for this gene, we analyzed the *C. albicans* ortholog *C7_00260C* using the CalCEN Co-expression network (29). GO term analysis revealed that 43 of 50 co-expressed genes fall under the “piecemeal microautophagy of the nucleus” term (Fig. S1). We also observed pseudohyphal filaments characterized by elongated cells with constricted separations between compartments in a mutant with an insertion in *CauELM1* (B9J08_002849), an ortholog of *S. cerevisiae ELM1* (YKL048C) (Fig. 2).

**Fig. 2.**
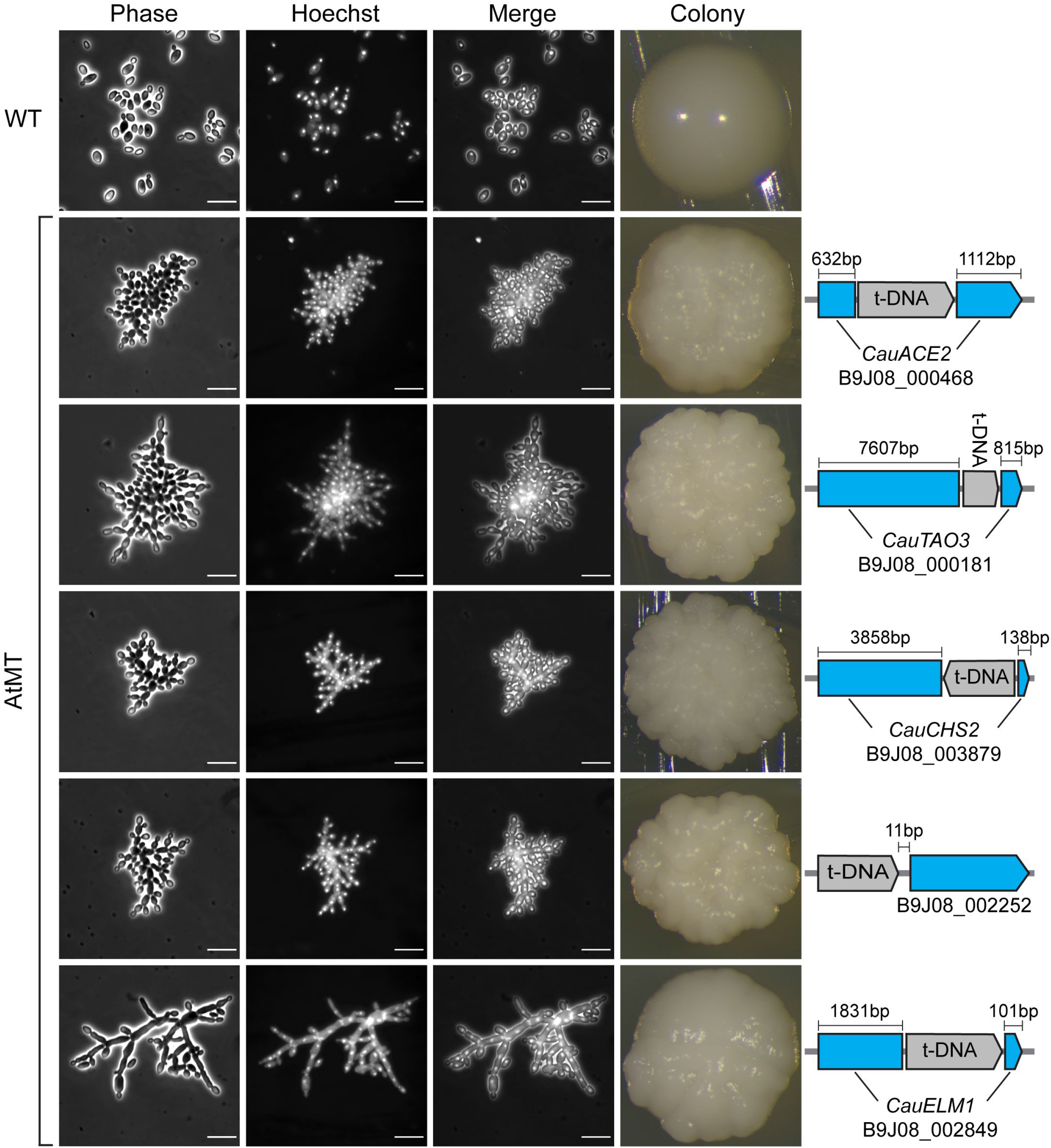
Transgene insertion sites associated with *C. auris* morphogenic mutants. Phase contrast, Hoescht 33342 staining, and colony morphologies demonstrate distinct morphogenic defects in five AtMT mutants (bottom) compared to wild-type *C. auris* AR382 (top). Identified transgene insertion sites were confirmed using Sanger sequencing (right). In all five cases, t-DNA insertion events were not accompanied by any additional insertions or deletions in the insertion locus. Scale bar = 10 μm.

A sixth insertional mutant identified by its irregular colony morphology exhibited elongated cell growth in short chains (Fig. S2). For this mutant, we identified T-DNA sequence both in the intergenic space upstream of the B9J08_002954 ORF and in the intergenic region upstream of the B9J08_002667 ORF from the B8441 reference sequence, but we were unable to amplify the complete insertion locus of either site from genomic DNA of the mutant. We hypothesize that a recombination event or other chromosomal rearrangement may have occurred following one or multiple T-DNA insertion events in this mutant, though further investigation is required to confirm this. Together, these findings identify key components in the regulation of cell separation in *C. auris*.

### Expression of Cas9 and sgRNA increases targeted integration in *C. auris*

To validate the insertional mutagenesis and confirm the role of identified genes in regulating the multicellular phenotypes we observed, it is important to be able to recapitulate the phenotype via clean deletions of the target genes. However, targeted homologous recombination has low efficiency in *C. auris*; in our hands, the rate of targeted integration events approaches or falls below 1% of all transformants for some isolates of *C. auris* when relying on homologous recombination alone (Fig 3D).

**Fig. 3.**
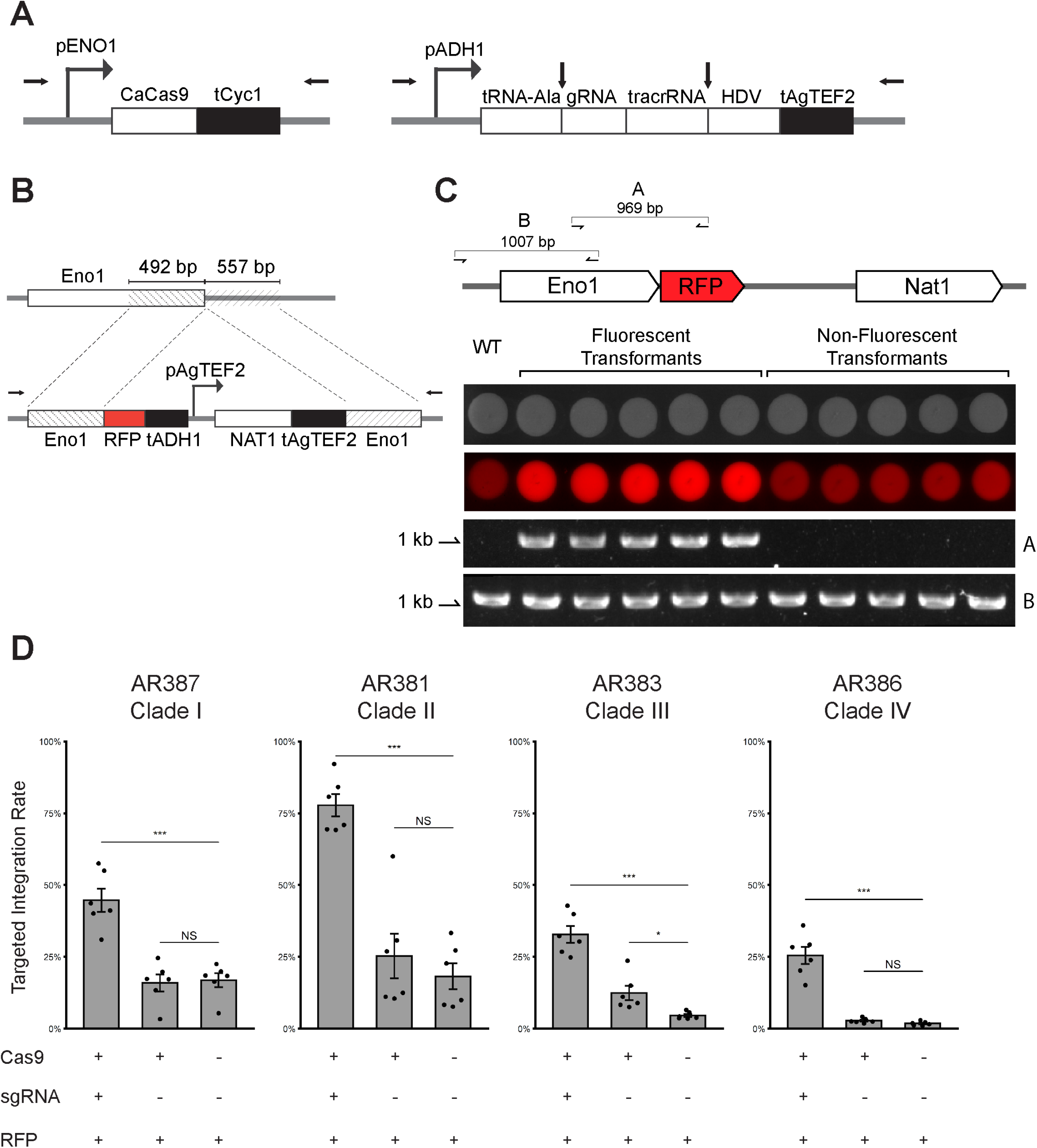
A CRISPR-Cas9 expression system promotes targeted transformation in four *C. auris* clades. (A) Structures of the Cas9 and sgRNA expression cassettes. *CAS9* is driven by the *C. auris ENO1* promoter and followed by the *CYC1* terminator. The sgRNA cassette is driven by the *C. auris ADH1* promoter and contains *C. auris* tRNA-Ala immediately upstream of the 20-bp gRNA sequence and hepatitis delta virus (HDV) ribozyme immediately downstream of the tracrRNA sequence. Predicted post-transcriptional cleavage sites are indicated by vertical arrows. Primer sites to generate linear transformation cassettes are indicated by horizontal arrows. (B) Design of the reporter cassette for measuring targeted integration. The cassette is flanked by approximately 500-bp homology to the *C. auris ENO1* C-terminus minus the stop codon and the region immediately downstream of *C. auris ENO1. RFP* and the *C. auris ADH1* terminator tag the *ENO1* gene at the C-terminus via a glycine linker to generate *ENO1-RFP* in targeted transformants. An independently-driven nourseothricin resistance cassette (NAT1) allows identification of total transformants, regardless of integration site, by selection with nourseothricin. (C) Targeted integration events are identifiable by colony fluorescence. Transformation of AR387 was performed using the reporter cassette described in panel B. Representative fluorescent transformants and non-fluorescent transformants were spotted onto YPD. Primer set A, spanning the *ENO1-RFP* junction, shows amplification only from fluorescent transformants. Primer set B, spanning a neighboring wild-type locus, shows amplification from all transformants and the wild type. (D) Expression of Cas9 and sgRNA promotes targeted integration rate. Transformation was performed in representative isolates from all four major *C. auris* clades with the linear transformation cassettes described in panels A and B. Transformations were performed with and without Cas9 and sgRNA elements; when absent, the cassettes were replaced with an equivalent volume of buffer. Targeted integration rate is expressed as the ratio of fluorescent colonies recovered to total nourseothricin resistant colonies recovered. Each point represents an individual transformation. Shown are the mean and standard error of the mean from three individual experiments, each performed in duplicate. *, *P* < 0.05; **, *P* < 0.01; ***, *P* < 0.001; NS, *P* > 0.05; Welch’s two sample t-test.

Transformation in *C. auris* can be facilitated by the use of Cas9 and sgRNA ribonucleoproteins; however, the DNA-based transient CRISPR-Cas9 expression approach used in *C. albicans* been previously shown to not increase efficacy for *C. auris* (7)personal communication, Sang Hu Kim). Recently, Ng and Dean reported variable increases in targeted transformation efficiency in *C. albicans* when using different promoters to drive the transcription of the sgRNA (30). We hypothesized that the low previous efficiency of the transient CRISPR system may be due to poor recognition of the *SNR52* promoter from *C. albicans*. Therefore, we developed a transient Cas9 and sgRNA expression system that can be used for efficient transformation in *C. auris* (19, 30). First, we generated expression cassettes for Cas9 and sgRNA using *C. auris*-specific promoters (Fig. 3A). We placed the CaCas9 cassette, which has been codon-optimized for expression in CTG clade fungi, under control of the *C. auris ENO1* promoter and the sgRNA cassette under control of the *C. auris ADH1* promoter. However, use of an RNA Polymerase II promoter would generate a transcript with a 5’ cap and 3’ polyA tail, ultimately detrimental to the gRNA targeting efficiency; therefore, to generate an sgRNA transcript with clean 5’ and 3’ ends after transcription, we included the *C. auris tRNA-Ala* sequence immediately upstream of the sgRNA and the hepatitis delta virus (HDV) Ribozyme sequence immediately downstream of the sgRNA. With this design, we anticipated cleavage at the 3’ end of the tRNA sequence by endogenous RNase A and self-catalyzed cleavage at the 5’ end of the HDV ribozyme (31, 32).

To assess the functional capacity of the Cas9 and sgRNA expression system to increase the efficiency of targeted integration in *C. auris*, we designed a reporter cassette that would allow for rapid and specific identification of targeted integration events among transformants (Fig. 3B). The reporter cassette contained approximately 500 bp homology to the C-terminus of *C. auris ENO1* and genomic sequence immediately downstream of *ENO1*. We removed the stop codon from the *ENO1* C-terminus homologous sequence and fused *RFP* to the *ENO1* C-terminus with a glycine linker. Because the *RFP* gene had no promoter element, we anticipated transformants would only demonstrate robust fluorescence if the reporter cassette integrated precisely in frame to tag the Eno1 protein and be driven by the endogenous *ENO1* promoter. The reporter cassette also included an independently-driven nourseothricin resistance (NAT^R^) cassette to allow identification of the total transformant population by selection on nourseothricin, regardless of integration site. To confirm that the reporter cassette specifically identified targeted integration events, we designed a PCR primer set spanning the *ENO1-RFP* junction and a primer set spanning a region of the *ENO1* locus native to the wild type. We performed transformation with the reporter cassette and collected five representative transformants that exhibited robust fluorescence and five that did not. Amplification of the region spanning the *ENO1-RFP* junction was only exhibited by the fluorescent transformants and not by the wild type or non-fluorescent transformants, while amplification of the wild-type sequence was exhibited by all the transformants and the wild-type strain (Fig. 3C). This demonstrates that the ratio of fluorescent to non-fluorescent colonies is a reliable measure of targeted transformation efficiency.

We observed variable targeted transformation efficiency among *C. auris* isolates of different genetic backgrounds (Fig. 3D). We therefore sought to determine whether our Cas9 and sgRNA expression system promoted targeted transformation in multiple genetically diverse *C. auris* isolates. We performed transformations of *C. auris* isolates from all four major clades using the reporter cassette alone or in combination with the Cas9 and sgRNA expression cassettes (Fig. 3D). The targeted integration rate under each transformation condition was determined by dividing the number of fluorescent colonies by the total number of transformant colonies. For each isolate, transformation including both the Cas9 and sgRNA cassettes significantly increased targeted integration efficiency compared to transformation with the reporter cassette alone, though absolute rates of targeted integration varied between strains. The *ENO1* C-terminus homologous arm encoded by the reporter cassette showed 100% sequence identity in all four isolates, while AR381 and AR383 exhibited shared 4 nucleotide variants out of 557 bp in the downstream homologous arm and AR386 showed a single nucleotide variant in the same region (Fig. S3). Therefore, differences in the targeted integration efficiency could not be explained by differential homology to the reporter cassette. Taken together, these observations indicate the Cas9 and sgRNA expression cassettes successfully promote targeted transformation in all four *C. auris* clades.

### *CauACE2* and *CauELMI* are regulators of *C. auris* morphogenesis

Using these tools, we were able to investigate the function of the genes implicated in *C. auris* morphogenic regulation by AtMT. Deletion of *ACE2* in AR382 (Clade I) resulted in constitutively aggregating cells with individual cells connected at septa, suggestive of a failure of budding daughter cells to separate from mother cells (Fig. 4A). Mutations in *ACE2* in *S. cerevisiae* or in *C. albicans* result in an aggregating, multicellular phenotype similar to that exhibited by *C. auris ACE2* mutants, suggesting *C. auris* has maintained conservation of *ACE2* in regulating morphogenesis (33–35). Deletion of *ELM1* resulted in filamentous pseudohyphal growth with constrictions at septa and numerous highly vacuolar cell compartments (Fig. 4A). *S. cerevisiae* strains with mutations in *ELM1* exhibit similar polarized, elongated growth phenotypes (36). In *C. glabrata*, mutation of *ELM1* results in elongated cells but does not fully recapitulate the pseudohyphal morphology exhibited by *S. cerevisiae* or by the *C. auris Δelm1* mutant (37). Similar phenotypes were observed for *Δace2* and *Δelm1* mutants in AR381 (Clade II), suggesting conserved roles of these regulators across *C. auris* clades (Fig. 4B).

**Fig. 4.**
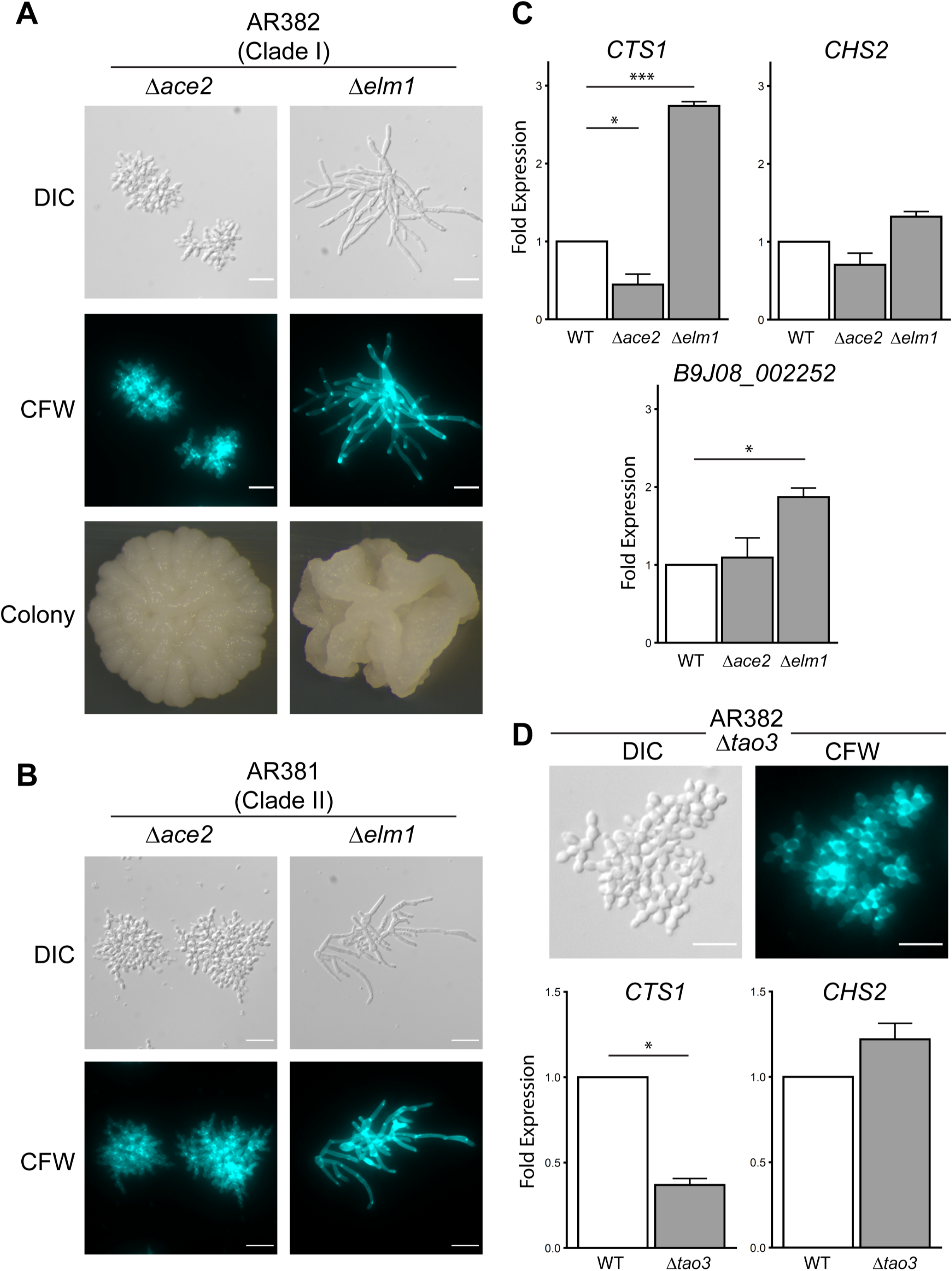
*ACE2* and *ELM1* are regulators of *C. auris* morphogenesis. (A) Microscopy of *Δace2* and *Δelm1* strains in the AR382 (Clade I) genetic background. Representative images shown for DIC, cells stained with calcofluor white, and colonies formed on YPD agar. (B) *ACE2* and *ELM1* regulate morphogenesis across *C. auris* clades. Microscopy of *Δace2* and *Δelm1* strains in the AR381 (Clade II) genetic background. Representative images shown for DIC and cells stained with calcofluor white. (C) *ACE2* and *ELM1* regulate putative chitinase *CTS1* transcription. Wild-type (AR382), *Δace2,* and *Δelm1* cells were grown to exponential phase in YPD at 30 °C prior to RNA extraction and RT-qPCR analysis of upregulated and downregulated genes. (D) Microscopy of *Δtao3* in the AR382 (Clade I) genetic background. Representative images shown for DIC and cells stained with calcofluor white. For RT-qPCR, wild-type (AR382) and *Δtao3* cells were grown to exponential phase in YPD at 30 °C prior to RNA extraction. Scale bar = 20 μm for all microscopy images. RT-qPCR results are presented as fold expression relative to the wild-type strain, normalized to *ACT1* expression. Data represent mean and standard error of the mean from three replicates. Statistically significant fold changes are indicated; *, *P* < 0.05; **, *P* < 0.01; ***, *P* < 0.001; Welch’s two sample t-test.

To test whether the regulation of cell wall maintenance genes by these regulatory pathways was also conserved between organisms, we investigated transcriptional changes associated with the mutants (Fig. 4C). The *Δace2* mutant exhibited decreased expression of the putative chitinase *CauCTS1* (B9J08_002761), consistent with the role of *Sc*Ace2 in regulating *ScCTS1* to promote degradation of the primary septum during daughter cell separation (38). While *Δelm1* cells also remained septally conjoined, the *Δelm1* mutant exhibited increased expression of *CauCTS1*. The *Δelm1* mutant also exhibited a modest increase in the expression of *B9J08_002252*, the gene of unknown function identified in our insertional mutagenesis to be associated with aggregating morphology. Disruption of the putative chitin synthase gene *CauCHS2* was also associated with an aggregating cell morphology in our AtMT screen. Orthologs of *CHS2* in *S. cerevisiae* or *C. albicans* are thought to catalyze the formation of primary septum chitin; defects in these orthologous genes result in multicellular clumps or chains with abnormal cytokinesis patterns (39, 40). We observed little change in the expression of *CauCHS2* in *Δace2* or *Δelm1* mutants (Fig. 4C), suggesting chitin synthase transcriptional regulation in *C. auris* is not altered in response to perturbations in chitinase expression. Deletion of *TAO3*, another gene associated with aggregating cell morphology in our AtMT screen, resulted in aggregating cells similar to *Δace2* (Fig. 4D). In *S. cerevisiae*, Tao3 associates with kinases Kic1 and Cbk1 as part of the Regulation of *ACE2* Morphogenesis (RAM) pathway. Phosphorylation of Ace2 by Cbk1 results in its accumulation in daughter cell nuclei, where it regulates the expression of enzymes that mediate septum degradation (41). Consistent with this role, *CTS1* expression was significantly downregulated in *Δtao3* cells compared to wild type (Fig. 4D). We observed no change in the expression of *CHS2* in the *Δtao3* mutant, suggesting *CHS2* transcriptional regulation is not controlled by *TAO3*-dependent components of the RAM pathway (Fig. 4D). Taken together, these observations identify *ACE2* and *ELM1* as key regulators of *C. auris* morphogenesis associated with transcriptional regulation of *CTS1*.

## Discussion

We have developed new approaches to performing facile, cost-effective forward and reverse genetic manipulation in *C. auris*. Using these tools, we identified functional conservation of chitinase and chitin synthase regulatory pathways, disruption of which results in aggregating, multicellular growth in *C. auris*. Cell wall chitin remodeling during growth and cell separation involves parallel expression of both chitinase and chitin synthase activities (42). Direct associations to the regulation of chitinase and chitin synthase were especially striking considering multiple genes involved in these processes were identified from a low-saturation library of approximately 2000 mutants. We also uncovered a novel *C. auris* pseudohyphal mutant, *Δelm1*, demonstrating the ability of *C. auris* to sustain filamentous growth. Our work represents part of a growing global effort to understand the biology of this emerging pathogen by offering alternative methods of improving its genetic tractability. We demonstrated the ability of a *C. auris* CRISPR-Cas9 expression system to consistently and significantly improve targeted integration of a transformation cassette in representative isolates from all four major *C. auris* clades. Targeted integration rates were increased to levels at which mutants of interest can readily be identified by PCR or phenotypic screening. While this level of efficiency was associated with approximately 500 bp arms of homology, we successfully performed deletion of *CauTAO3* using a transformation cassette with only 50-70 bp of homology, albeit with reduced targeted transformation efficiency. Our work, in concert with similar advancements such as successful resistance marker recycling in *C. auris* (18, 43), will promote improved accessibility to mechanistic understanding of the genetic machinery in *C. auris*.

From our work, we identified *CauACE2* to be a key regulator of morphogenesis. In *S. cerevisiae, ACE2* daughter cell nuclear localization is regulated by the RAM pathway Kic1-Cbk1 kinase complex (41). *ScTAO3*, sometimes called *PAG1*, physically associates with both *Sc*Kic1 and *Sc*Cbk1 and may mediate activation of Cbk1 by Kic1 (44, 45). Disruption of *ScTAO3* or downstream *ScACE2* results in cell aggregates and a failure of daughter cells to separate from mother cells during budding (38, 44, 45). We observed similar aggregating phenotypes in *Δace2* and *Δtao3* mutants in *C. auris*. We therefore propose functional conservation of *ACE2* and the RAM regulatory pathway in *C. auris* (Fig. 5). Downstream of this pathway, we identified a putative chitinase, *CauCTS1* (B9J08_002761), that was downregulated in *ΔCauace2* compared to the wild type. The sequence of *CauCTS1* contains no GPI-anchor signal, and so is likely more closely related functionally to the secreted chitinases *ScCTS1* in *S. cerevisiae* and its functional homolog *CaCHT3* in *C. albicans* than to *CaCHT2* in *C. albicans* (46). The regulation of *CauCTS1* by *CauACE2* is consistent with homologous pathways in *S. cerevisiae* and *C. albicans*, in which chitin degradation in the primary septum is mediated by the *ACE2*-regulated ScCts1 or CaCht3 proteins (34, 47). Interestingly, an experiment performing laboratory evolution of *S. cerevisiae* in a bioreactor resulted in multicellular, fast-sedimenting strains that were associated with mutations in *ACE2* (33). The design of the bioreactors in this example may have provided a selective advantage for multicellular growth due to increased sedimentation rate of cell aggregates compared to planktonic cells. An environmental niche may exist that produces a similar selective pressure against the regulatory network upstream of *CTS1* by offering a selective advantage for aggregating cells. Constitutively aggregating strains of *C. auris* have been isolated from clinical samples (12–15). If an environmental reservoir for *C. auris* is aquatic in nature, as some hypotheses suggest (48–50), a fast-sedimenting aggregative phenotype may confer a selective advantage by offering increased nutritional access through sedimentation or resistance to dispersal by moving water. It is tempting to speculate whether aggregating *C. auris* isolates have evolved a multicellular phenotype through a similar selective pressure on chitinase or chitin synthase regulation prior to introduction to a human host. Further characterization of the environmental reservoirs for *C. auris* may offer insight regarding the selective pressures driving similar phenotypes.

**Fig. 5.**
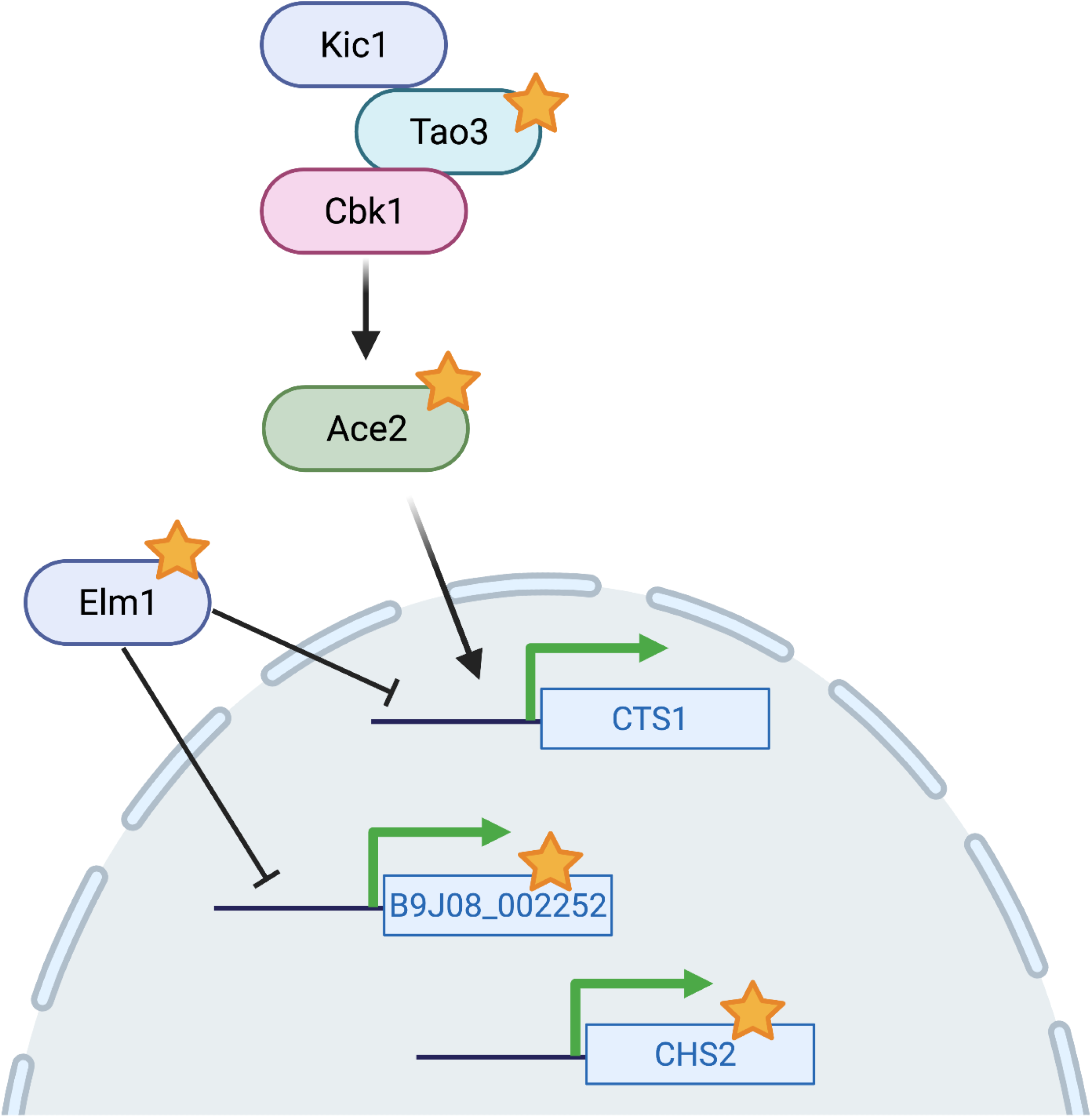
Proposed model for the regulation of morphogenesis in *C. auris*. Ace2 mediates cell separation through transcriptional upregulation of the putative chitinase *CTS1*. Conserved components of the RAM pathway Kic1 and Cbk1 interact through Tao3 to promote the transcription factor activity of Ace2. Elm1 is a negative regulator of *CTS1;* cells defective in Elm1 activity exhibit aberrant pseudohyphal morphology. Elm1 also serves as a negative regulator of *B9J08_002252,* which is associated with cell separation through unknown mechanisms. Genes identified as associated with aggregating or filamentous morphology through AtMT are indicated with stars.

We also observed aggregating growth in an insertional mutant in a putative chitin synthase, *CauCHS2* (B9J08_003879). The orthologous gene *ScCHS2* is required for formation of the primary septum in budding yeast and its disruption results in abnormal, multicellular clusters (39). Careful regulation of both the chitin synthase, catalyzing the formation of the primary septum, and the chitinases that degrade the primary septum is critical in successful cell separation. A “unitary model” of cell wall growth suggests coordinated regulation of chitin synthesis and degradation (51). In contrast to this model, our observations that expression of *CauCHS2* remained relatively constant in *Δace2* or *Δelm1 C. auris* strains, which exhibited decreased or elevated *CTS1* expression respectively, indicate regulation of *CHS2* independent of *CTS1*, consistent with a report that regulation of chitin synthase and chitinase activities are independent in *C. albicans* and *S. cerevisiae* (52). Still, the temporal regulation of the contrasting activities of *CTS1* and *CHS2* must be essential for effective cell separation. In support of this notion, recent CHIP-exo data indicates that *ACE2* and *CHS2* are both targets of the forkhead transcription factor Fkh2 in *S. cerevisiae*, suggesting Fkh2 may act as a regulatory hub governing the contrasting chitin synthase and chitinase activities during cell separation (53). While the RAM pathway regulatory kinase Cbk1 targets Fkh2 in addition to Ace2 in *C. albicans*, our findings indicate such an interaction does not directly transcriptionally regulate *CHS2* expression in *C. auris* in a Tao3-dependent manner (54, 55). Future work may yet reveal insights into the concerted regulation of chitin synthase and chitinase activity during cell separation. Our findings suggest the roles of the RAM pathway and *ACE2* in regulating *CTS1* are conserved in *C. auris* and critical for morphogenesis.

While the role of the serine-threonine kinase *ELM1* in regulating polar bud growth and morphogenic differentiation in *S. cerevisiae* has been long understood, its role in pathogenic fungi is largely unexplored (36, 56). One report demonstrated that deletion of *CgELM1* in *C. glabrata* results in moderately elongated cell growth, though this strain fails to recapitulate the fully pseudohyphal phenotype exhibited by *S. cerevisiae* or *C. auris* (37). We observed elongated cells growing in pseudohyphal chains associated with an insertion event near the C-terminus of *CauELM1*. However, the full *Δelm1 C. auris* strains exhibited a slightly different, more filamentous cell morphology. In *S. cerevisiae*, deletion of the C-terminal domain of *ELM1* results in increased Elm1 kinase activity, suggesting this domain may have autoinhibitory function (57). This phenotype is associated with pseudohyphal growth with a cell morphology distinct from that demonstrated by *ΔScelm1* (57). The distinct but similarly pseudohyphal phenotypes associated with disruption of the *C. auris ELM1* C-terminus and *ΔCauelm1* suggests similar *ELM1* regulation may exist in *C. auris*. Intriguingly, the pseudohyphal *Δelm1 C. auris* mutant exhibited a significant increase in the expression of *CTS1* compared to the wild type. This is in contrast to *Δelm1* in *C. glabrata*, which exhibited decreased expression of *CgCTS1* compared to wild type (37). Further characterization of Elm1 in diverse fungal species may yet reveal substantial variation in its function. The role that increased *CTS1* expression in *ΔCauelm1* plays in contributing to pseudohyphal growth is unclear. One report indicated reduced expression of the *CTS1* homolog *CaCHT3* in hyphal *C. albicans* compared to *C. albicans* grown in the yeast form (58). However, total chitinase activity was increased in *C. albicans* hyphae compared to yeast (52). Whether *C. auris* pseudohyphal growth is controlled by a similar chitinase function as *C. albicans* hyphal growth remains to be determined.

In sum, our work demonstrates an accessible approach to genetic engineering of *C. auris*, facilitating further understanding of the biology of this emerging pathogen. Using new forward and reverse genetic approaches, we characterized conserved and divergent key regulators of morphogenesis in *C. auris*.

## Materials and Methods

### Strains and Growth Conditions

A list of *C. auris* and *A. tumefaciens* strains used in this study are listed in Table S1. Unless specified otherwise, *C. auris* cells were grown at 30 °C in YPD liquid media (1% yeast extract, 2% peptone, 2% dextrose) with constant agitation. All strains were maintained in frozen stocks of 25% glycerol at −80 °C.

### Plasmids

A list of all plasmids used in this study is included in Table S2.

A list of all primers used in this study is included in Table S3.

#### pTO128

An *Agrobacterium* Ti-plamid was constructed to include the CaNAT1 nourseothricin resistance cassette (59) in the pPZP-NEO1 backbone (60). The CaNAT1 cassette was excised at the SacI and SalI restriction sites from pLC49 (61) and ligated between the SacI and SalI restriction sites of pPZP-NEO1, replacing the neomycin resistance cassette with CaNAT1 to form pTO128 (pPZP-NATca). pTO128 was subsequently electroporated into *A. tumefaciens* strain EHA 105 (62) using a Bio-Rad MicroPulser Electroporator.

#### pTO135

A cassette for expression of Cas9 was maintained in the pUC19 cloning vector backbone (63). To form pTO135 (pCauCas9), the Cas9 expression cassette was PCR amplified from pLC963 (64) without the promoter sequence using primers oTO114-oTO115. A promoter region consisting of 1000 bp upstream of the *C. auris ENO1* gene (B9J08_000274) was PCR amplified from genomic DNA isolated from *C. auris* strain AR387 using primers oTO112-oTO113. The pUC19 vector backbone was amplified using primers oTO116-oTO117. The promoter sequence and Cas9 expression cassette were assembled into the multiple cloning site of pUC19 using the NEBuilder HIFI DNA Assembly master mix (NEB #E2621) according to the manufacturer’s instructions.

#### pTO136

A cassette for expression of sgRNA was maintained in the pUC19 cloning vector backbone (63). To form pTO136 (pCausgRNA), a promoter region consisting of 901 bp upstream of the *C. auris ADH1* gene (B9J08_004331) was PCR amplified using primers oTO118-oTO119 and assembled along with a synthesized DNA fragment (Genscript, Piscataway, NJ, USA) containing sequence from *C. auris* tRNA-Ala (B9J08_003096), a 20-bp gRNA sequence targeting the *ENO1* locus, a tracrRNA sequence, and an HDV ribozyme (30) into the multiple cloning site of pUC19 using the NEBuilder HIFI DNA Assembly master mix according to the manufacturer’s instructions. The pUC19 vector backbone was amplified using primers oTO120-oTO121.

#### pTO137

A reporter cassette for tagging the *C. auris ENO1* gene with RFP was maintained in the pUC19 cloning vector backbone. To form pTO137, the *RFP* construct was PCR amplified from pLC1047 (65) using primers oTO124-oTO125; a terminator sequence consisting of 933 bp downstream of the *C. auris ADH1* gene was PCR amplified from genomic DNA isolated from *C. auris* strain AR387 using primers oTO126-oTO127; the CaNAT1 expression cassette including *TEF* promoter and terminator sequence was amplified from pLC49 using primers oTO128-oTO129; flanking regions containing homology to 492 bp at the C-terminal end of the *C. auris ENO1* gene minus the stop codon and 557 bp immediately 3’ of the *C. auris ENO1* gene were amplified from genomic DNA isolated from *C. auris* strain AR387 using primers oTO122-oTO123 and oTO130-oTO131 respectively; the pUC19 vector backbone was amplified using primers oTO132-oTO133. Fragments were assembled into the multiple cloning site of pUC19 using the NEBuilder HIFI DNA Assembly master mix according to the manufacturer’s instructions.

#### pTO154

A repair cassette for deleting *ELM1* with *CaNAT1* was maintained in the pUC19 cloning vector backbone. To form pTO154 (*pELM1::NAT*), 501 bp immediately 5’ of *ELM1* (B9J08_002849) and 502 bp immediately 3’ of *ELM1* were amplified from AR387 genomic DNA using primers oTO317-oTO318 and oTO321-oTO337 respectively; the *CaNAT1* expression cassette was amplified from pLC49 using primers oTO319-oTO320; the pUC19 vector backbone was amplified using primers oTO323-oTO324. Fragments were assembled into the multiple cloning site of pUC19 using the NEBuilder HIFI DNA Assembly master mix according to the manufacturer’s instructions.

#### pTO155

A repair cassette for deleting *ACE2* with *CaNAT1* was maintained in the pUC19 cloning vector backbone. To form pTO155 (*pACE2::NAT*), 500 bp immediately 5’ of *ACE2* (B9J08_000468) and 498 bp immediately 3’ of *ACE2* were amplified from AR387 genomic DNA using primers oTO325-oTO326 and oTO329-oTO330 respectively; the *CaNAT1* expression cassette was amplified from pLC49 using primers oTO327-oTO328; the pUC19 vector backbone was amplified using primers oTO331-oTO332. Fragments were assembled into the multiple cloning site of pUC19 using the NEBuilder HIFI DNA Assembly master mix according to the manufacturer’s instructions.

All *C. auris* genomic sequence data was obtained from the *C. auris* B8441 reference genome on fungidb.org (66). All plasmid assemblies were verified by restriction digest and sanger sequencing.

### *Agrobacterium tumefaciens-Mediated* Transformation (AtMT)

AtMT was performed as previously described with minor modifications (67). Briefly, *A. tumefaciens* strain EHA 105 harboring the pTO128 (pPZP-NATca) plasmid was cultured overnight at 30 °C in liquid Luria-Bertani (LB) media containing kanamycin. *A. tumefaciens* cells were harvested by centrifugation, washed once with sterile, ultrapure water, then resuspended at a final OD_600_ of 0.15 in liquid Induction Medium (IM) supplemented with 100 μM Acetosyringone 3’,5’-dimethoxy-4-hydroxyacetophenone (AS) (27) and incubated at room temperature for 6 h with constant shaking. Recipient *C. auris* AR382 cells were harvested from an overnight culture grown at 30 °C in YPD by centrifugation then resuspended in sterile, ultrapure water at a final OD_600_ of 1.0. To determine the maximal transformation efficiency, prepared cells were combined at *C. auris:A. tumefaciens* ratios (v/v) of 1:1, 1:3, and 3:1. The combined cultures were incubated on IM with AS agar at 30 °C for 2, 4, and 7 days. Cells were then harvested into liquid YPD, washed three times with fresh YPD, then spread-plated onto YPD agar containing 200 μg/mL nourseothricin and 200 μg/mL cefotaxime and incubated at 30 °C for 2 days. Transformation efficiency was determined by dividing the total number of recovered *C. auris* CFU by the total input number of *C. auris* cells for each condition. To screen for *C. auris* morphologic mutants, *A. tumefaciens* and *C. auris* cells were prepared as described above then equal volumes combined and the mixed culture was incubated on IM with AS agar at 30 °C for 4 days. Cells were then harvested into liquid YPD, washed three times with fresh YPD, then spread-plated onto YPD agar containing 200 μg/mL nourseothricin and 200 μg/mL cefotaxime. Plates were incubated at 30 °C for 2 days. Colonies were then screened visually for those exhibiting a wrinkled colony morphology.

### Genomic DNA Isolation

Genomic DNA was isolated from *C. auris* morphological mutants to be used for downstream sequencing and insertion site mapping using a phenol-chloroform extraction. Briefly, cells were incubated overnight at 30 °C in liquid YPD then harvested by centrifugation and resuspended in breaking buffer (2% (v/v) Triton X-100, 1% (w/v) SDS, 100 mM NaCl, 10mM Tris-Cl, 1mM EDTA). DNA was extracted by bead beating into PCA then extracted into Chloroform. Following precipitation by ethanol, extracted DNA was resuspended in TE buffer and treated with RNase A. Genomic DNA quality was assessed by 1% agarose gel electrophoresis.

### ATMT Transgene Mapping

Mapping of tDNA insertion sites was performed similarly to methods previously described (28). Genomic DNA isolated from six morphogenic mutants was collected and pooled into two pools, each containing equal amounts by mass of genomic DNA from three individual mutants. Library preparation, quality control and Whole Genome Sequencing were performed by Microbial Genome Sequencing Center (MIGS, Pittsburg, PA, USA). Library preparation was performed based on the Illumina Nextera kit and sequencing performed on the Nextseq 550 platform generating 150 bp paired end reads for each pool. Sequencing data was analyzed using the Galaxy web platform public server at *usegalaxy.org* (68). Read quality was assessed using FASTQC and reads were trimmed using CutAdapt (69) with a Phred quality cutoff of 20. A linearized vector reference sequence of pTO121 (pPZP-CaNat) was generated from the circular vector sequence and 150 bp of sequence from the opposite border was added to each border of the linearized sequence. Reads were mapped to the linear pTO121 (pPZP-CaNAT) reference sequence using the Burrows-Wheeler Aligner with maximum exact matches (BWA-MEM) configured with default parameters except for minimum seed length = 50 and band width = 2 (70). Mapped reads were sorted based on position and sequences that extended beyond the left and right boundaries of the tDNA was extracted. The extracted sequences for each pool were aligned using Clustal Omega multiple sequence alignment (71) to identify consensus sequences for all independent insertion events within each pool. Consensus sequences were then mapped to the *C. auris* B8441 reference genome (GCA_002759435.2) using NCBI Blast. Primers specific to each identified insertion site were designed: oTO310 and oTO340 for *B9J08_002252*, oTO311 and oTO344 for *B9J08_003879*, oTO312 and oTO342 for *B9J08_002849*, oTO313 and oTO338 for *B9J08_000181*, oTO314 and oTO339 for *B9J08_000468,* oTO315 and oTO341 for *B9J08_002667,* and oTO316 and oTO343 for *B9J08_002954*. These were used to amplify the identified insertion regions in conjunction with insertion-specific primers oTO6 and oTO90 using the genomic DNA from each of the six mutants as templates. Individual insertions were attributed to individual mutants based on amplicon length. Amplicons containing tDNA insertions were Sanger Sequenced to generate insertion maps for each mutant.

### *C. auris* Transformation

Transformation of *C. auris* was performed as described previously, with minor modifications (17). To generate *ENO1-RFP* strains, linear transformation cassettes encoding Cas9, sgRNA, and the RFP repair cassette were PCR amplified from pTO135, pTO136, and pTO137, respectively, using primers oTO18-oTO19. To generate *Δelm1* and *Δace2* strains, a linear Cas9 cassette was amplified from pTO135 using primers oTO18-oTO19, linear repair cassettes were amplified from pTO154 for *ELM1::NAT* using primers oTO18-oTO19 and pTO155 for *ACE2::NAT* using primers oTO18-oTO19. To generate *Δtao3*, a linear repair cassette incorporating 50-70 bp homology to either end of the target gene flanking the NAT cassette was amplified from pTO137 using primers oTO353-oTO354. Linear sgRNA cassettes were amplified from pTO136 using fusion PCR as described previously to replace the gRNA sequence with gRNA targeting each gene for deletion (19). Fusion fragments were amplified using primers oTO333-oTO225 and oTO224-oTO334 to target *ELM1,* oTO335-oTO225 and oTO224-oTO336 to target *ACE2,* and oTO356-oTO224 and oTO355-oTO225 to target *TAO3*. Each pair of fragments with overlapping sequences were spliced on extension using oTO18-oTO19. PCR products were purified with a Zymo DNA Clean & Concentrator kit (Cat no. D4034, Zymo Research) according to the manufacturer’s instructions. *C. auris* cells were incubated overnight at 30 °C in YPD to exponential phase, not exceeding OD_600_ of 2.2. Cells were harvested by centrifugation and resuspended in TE buffer with 100 mM Lithium Acetate then incubated with constant shaking at 30 °C for 1 h. DTT was added to the cells at a final concentration of 25 mM and incubation was continued for 30 min at 30 °C with constant shaking. The cells were harvested by centrifugation; washed once with ice-cold, sterile, ultrapure water; washed once with ice-cold 1 M Sorbitol; then resuspended in ice-cold 1 M Sorbitol. 40 μL of competent cells were added to a pre-chilled 2 mm-gap electro-cuvette along with 1 μg each of the PCR amplified linear transformation cassettes encoding Cas9, sgRNA, and the repair cassette. Alternatively, to compare targeted integration efficiency, an equal volume of Zymo elution buffer was added instead of Cas9 or sgRNA cassettes. Cells were electroporated using a Bio-Rad MicroPulser Electroporator set to the programmed *P. pastoris* (PIC) protocol (2.0 kV, 1 pulse), recovered in 1 M Sorbitol, then resuspended in YPD and allowed 2 hrs of outgrowth at 30 °C with shaking. The cells were then spread-plated on YPD with 200 ug/mL nourseothricin and incubated at 30 °C.

To estimate the efficiency of targeted RFP integration among transformant colonies, transformation plates were imaged using a Typhoon FLA 9500 Bioimager fitted with a 532 nm filter. Fluorescent images were visualized using Fiji Software (72). An intensity threshold was set to identify transformant colonies exhibiting fluorescence. Five representative fluorescent colonies and five representative non-fluorescent colonies from transformations performed in AR387 were spotted onto YPD agar and grown at 30 °C for 2 days. A sample of the colony growth was collected from each colony and suspended in 15 uL water. An aliquot of this suspension was used as a template in PCR reactions with primers overlapping the junction of the predicted *ENO1-RFP* insertion site or a genomic region upstream of the junction present in the wild-type locus. Colony PCR was performed using Phire Plant Direct PCR Master Mix (F160S; Thermo Fisher Scientific) according to the manufacturer’s instructions. The proportion of transformant colonies with targeted integration was determined by dividing the number of colonies exhibiting fluorescence by the total number of transformant colonies.

### Fixed Cell Microscopy

*C. auris* cells were grown overnight at 30 °C in YPD. Cells were harvested by centrifugation for 1 min at 4000 rpm (1500 x g) and resuspended in methanol. The fixed cells were pelleted by centrifugation for 1 min at 4000 rpm then resuspended in PBS with 10 μg/mL Hoechst 33342 (Cayman Chemical; Item no. 15542). Aliquots of stained cells were loaded onto glass microscope slides and visualized using an Olympus IX70 Epifluorescent Microscope fitted with a Hamamatsu C11440 camera.

### Live Cell Microscopy

Cells were grown to mid-exponential phase at 30 °C in YPD and pelleted by centrifugation for 1 min at 4000 rpm (1500 x g) then resuspended in PBS. 5 μL cell suspension was combined with 1 μL of 0.1 g/L Calcofluor White stain and applied to a glass microscope slide. Stained cells were visualized using an Olympus IX70 Epifluorescent Microscope fitted with a Hamamatsu C11440 camera.

### Stereomicroscopy

*C. auris* cells were grown on YPD agar at 30 °C for 2-7 days to form colonies. Colonies were visualized using a Leica KL300 LED stereomicroscope.

### RNA Extraction

RNA extraction was performed as described previously (73). Briefly, cells were grown to mid-exponential phase at 30 °C in YPD and harvested by centrifugation. Cells were washed in PBS, then centrifuged and all liquid removed. Dry cell pellets were frozen on dry ice then stored at −80 °C overnight. Cell pellets were thawed and resuspended in 100 μL FE Buffer (98% formamide, 0.01M EDTA) at room temperature. 50 μL of 500 μm RNAse-free glass beads was added and the cell suspension was ground in 3 cycles of 30 sec using a BioSpec Mini-Beadbeater-16 (Biospec Products Inc., Bartlesville, OK, USA). The cell lysate was centrifuged to remove cell debris and the crude RNA extract collected from the supernatant. The extract was DNAse-treated and purified using a Qiagen RNeasy mini kit (Qiagen; 74104) as per the manufacturer’s instructions. RNA integrity was confirmed through agarose gel electrophoresis using the bleach gel method (74).

### RT-qPCR

cDNA was synthesized from isolated RNA using the AffinityScript qPCR cDNA Synthesis Kit (Agilent Technologies, 600559) according to the manufacturer’s instructions and used as a template for qPCR. qPCR was performed in triplicate using a BioRad CFXConnect Real Time System. Fold changes were calculated using the double-delta CT method with expression normalized to that of *ACT1* and compared to wild type. Amplification was measured for *ACT1* using primers oTO359-oTO360, for *CHS2* using primers oTO361-oTO362, for *CTS1* using primers oTO363-oTO364, for *B9J08_002252* using primers oTO365-oTO366, and for *ACE2* using primers oTO373-oTO374. The qRT-PCR was performed in triplicate for two biological replicates. Raw qPCR data can be found in Data S1.

### Co-Expression

The *C. albicans* ortholog of *B9J08_002252* was identified through orthology on the Candida Genome Database as C7_00260C_A. This was used as a query in CalCEN and the top 50 most co-expressed neighbors were identified. This set was then examined for putative function through GO term enrichment in the Candida Genome database. The network was visualized using Cytoscape.

## Data Availability

Data from Illumina sequences used to identify transgene insertion sites are available in the NCBI SRA under BioProject accession number PRJNA722500.

## Acknowledgements

We thank J. Andrew Alspaugh (Duke University) for generously donating *A. tumefaciens* EHA105 and pPZP-NEO. We also express our appreciation to the CDC for making *C. auris* isolates used in this study available through the Antibiotic Resistance Bank program.

D.J.S. was supported by NIH T32AI007528. T.R.O was supported by NIH KAI137299 (NIAID)

We declare that we have no conflicts of interest.

---

**Fig. S1** A *C. albicans* ortholog of *B9J08_002252* is coexpressed with genes involved in piecemeal autophagy of the nucleus. For the *C. albicans* gene *C7_00260C,* a putative ortholog of the *C. auris* gene *B9J08_002252,* coexpressed genes were identified and analyzed for GO term association using the CalCEN coexpression network. Each node represents an individual gene and each edge corresponds to the relative degree of coexpression. 43 of 50 coexpressed genes fall under the “Piecemeal autophagy of the nucleus” GO term (dark green) and 7 fall under “GO term unknown, no annotation available” (light blue).

**Fig. S2** Phase contrast and Hoechst 33342 stain imaging of a sixth insertional mutant identified with irregular morphology through AtMT. The exact genomic locations of transgene insertion sites in this mutant could not be determined.

**Fig. S3** *C. auris* isolates exhibit differential homology to the *ENO1* 3’ homologous arm used in the targeted transformation efficiency reporter cassette. The sequence of the 3’ homologous arm used in the cassette is provided. A pairwise alignment between this sequence and the genomic sequence corresponding to the homologous region in each of the four isolates tested for targeted transformation efficiency (AR387, AR381, AR383, AR386) indicates differences in homology to the reporter cassette. Homology at any given position is indicated by ‘.’ while a nucleotide polymorphism at any given position is indicated by A, T, C, or G.

**Table S1** Strains used in this study.

**Table S2** Plasmids used in this study.

**Table S3** Oligonucleotides used in this study.

**Data S1** Raw data for qPCR in Figure 4

## References

1. Satoh K, Makimura K, Hasumi Y, Nishiyama Y, Uchida K, Yamaguchi H. 2009. *Candida auris* sp. nov., a novel ascomycetous yeast isolated from the external ear canal of an inpatient in a Japanese hospital. Microbiol Immunol 53:41–44.

2. Chakrabarti A, Sood P. 2021. On the emergence, spread and resistance of *Candida auris*: host, pathogen and environmental tipping points. J Med Microbiol

3. Chow NA, Muñoz JF, Gade L, Berkow EL, Li X, Welsh RM, Forsberg K, Lockhart SR, Adam R, Alanio A, Alastruey-Izquierdo A, Althawadi S, Araúz AB, Ben-Ami R, Bharat A, Calvo B, Desnos-Ollivier M, Escandón P, Gardam D, Gunturu R, Heath CH, Kurzai O, Martin R, Litvintseva AP, Cuomo CA. 2020. Tracing the Evolutionary History and Global Expansion of *Candida auris* Using Population Genomic Analyses. MBio 11.

4. Chow NA, de Groot T, Badali H, Abastabar M, Chiller TM, Meis JF. 2019. Potential Fifth Clade of *Candida auris*, Iran, 2018. Emerg Infect Dis 25:1780–1781.

5. Forgács L, Borman AM, Prépost E, Tóth Z, Kardos G, Kovács R, Szekely A, Nagy F, Kovacs I, Majoros L. 2020. Comparison of in vivo pathogenicity of four *Candida auris* clades in a neutropenic bloodstream infection murine model. Emerg Microbes Infect 9:1160–1169.

6. Huang X, Hurabielle C, Drummond RA, Bouladoux N, Desai JV, Sim CK, Belkaid Y, Lionakis MS, Segre JA. 2020. Murine model of colonization with fungal pathogen *Candida auris* to explore skin tropism, host risk factors and therapeutic strategies. Cell Host Microbe

7. Kim SH, Iyer KR, Pardeshi L, Muñoz JF, Robbins N, Cuomo CA, Wong KH, Cowen LE. 2019. Genetic Analysis of *Candida auris* Implicates Hsp90 in Morphogenesis and Azole Tolerance and Cdr1 in Azole Resistance. MBio 10.

8. Bravo Ruiz G, Ross ZK, Gow NAR, Lorenz A. 2020. Pseudohyphal Growth of the Emerging Pathogen *Candida auris* Is Triggered by Genotoxic Stress through the S Phase Checkpoint. mSphere 5.

9. Wang X, Bing J, Zheng Q, Zhang F, Liu J, Yue H, Tao L, Du H, Wang Y, Wang H, Huang G. 2018. The first isolate of *Candida auris* in China: clinical and biological aspects. Emerg Microbes Infect 7:93.

10. Yue H, Bing J, Zheng Q, Zhang Y, Hu T, Du H, Wang H, Huang G. 2018. Filamentation in *Candida auris*, an emerging fungal pathogen of humans: passage through the mammalian body induces a heritable phenotypic switch. Emerg Microbes Infect 7:188.

11. Fan S, Yue H, Zheng Q, Bing J, Tian S, Chen J, Ennis CL, Nobile CJ, Huang G, Du H. 2021. Filamentous growth is a general feature of *Candida auris* clinical isolates. Med Mycol

12. Borman AM, Szekely A, Johnson EM. 2016. Comparative Pathogenicity of United Kingdom Isolates of the Emerging Pathogen *Candida auris* and Other Key Pathogenic Candida Species. mSphere 1.

13. Short B, Brown J, Delaney C, Sherry L, Williams C, Ramage G, Kean R. 2019. *Candida auris* exhibits resilient biofilm characteristics in vitro: implications for environmental persistence. J Hosp Infect 103:92–96.

14. Szekely A, Borman AM, Johnson EM. 2019. Candida auris Isolates of the Southern Asian and South African Lineages Exhibit Different Phenotypic and Antifungal Susceptibility Profiles In Vitro. J Clin Microbiol 57.

15. Brown JL, Delaney C, Short B, Butcher MC, McKloud E, Williams C, Kean R, Ramage G. 2020. *Candida auris* Phenotypic Heterogeneity Determines Pathogenicity In Vitro. mSphere 5.

16. Sherry L, Ramage G, Kean R, Borman A, Johnson EM, Richardson MD, Rautemaa-Richardson R. 2017. Biofilm-Forming Capability of Highly Virulent, Multidrug-Resistant *Candida auris*. Emerg Infect Dis 23:328–331.

17. Grahl N, Demers EG, Crocker AW, Hogan DA. 2017. Use of RNA-Protein Complexes for Genome Editing in Non-albicans *Candida* Species. mSphere 2.

18. Rybak JM, Doorley LA, Nishimoto AT, Barker KS, Palmer GE, Rogers PD. 2019. Abrogation of Triazole Resistance upon Deletion of CDR1 in a Clinical Isolate of *Candida auris*. Antimicrob Agents Chemother 63.

19. Min K, Ichikawa Y, Woolford CA, Mitchell AP. 2016. *Candida albicans* Gene Deletion with a Transient CRISPR-Cas9 System. mSphere 1.

20. Fan Y, Lin X. 2018. Multiple Applications of a Transient CRISPR-Cas9 Coupled with Electroporation (TRACE) System in the *Cryptococcus neoformans* Species Complex. Genetics 208:1357–1372.

21. Soltani J, van Heusden GPH, Hooykaas PJJ. 2008. Agrobacterium-Mediated Transformation of Non-Plant Organisms, p. 649–675. In Tzfira, T, Citovsky, V (eds.), Agrobacterium: From Biology to Biotechnology. Springer New York, New York, NY.

22. Cleene MD, De Cleene M, De Ley J. 1976. The host range of crown gall. The Botanical Review.

23. Michielse CB, Hooykaas PJJ, van den Hondel CAMJJ, Ram AFJ. 2005. *Agrobacterium*-mediated transformation as a tool for functional genomics in fungi. Curr Genet 48:1–17.

24. Hooykaas PJJ, van Heusden GPH, Niu X, Reza Roushan M, Soltani J, Zhang X, van der Zaal BJ. 2018. *Agrobacterium-Mediated* Transformation of Yeast and Fungi. Curr Top Microbiol Immunol 418:349–374.

25. Lutgring JD, Machado M-J, Benahmed FH, Conville P, Shawar RM, Patel J, Brown AC. 2018. FDA-CDC Antimicrobial Resistance Isolate Bank: a Publicly Available Resource To Support Research, Development, and Regulatory Requirements. J Clin Microbiol 56.

26. McClelland CM, Chang YC, Kwon-Chung KJ. 2005. High frequency transformation of *Cryptococcus neoformans* and *Cryptococcus gattii* by *Agrobacterium tumefaciens*. Fungal Genet Biol 42:904–913.

27. Bundock P, den Dulk-Ras A, Beijersbergen A, Hooykaas PJ. 1995. Trans-kingdom T-DNA transfer from *Agrobacterium tumefaciens* to *Saccharomyces cerevisiae*. EMBO J 14:3206–3214.

28. Park D, Park S-H, Ban YW, Kim YS, Park K-C, Kim N-S, Kim J-K, Choi I-Y. 2017. A bioinformatics approach for identifying transgene insertion sites using whole genome sequencing data. BMC Biotechnol 17:67.

29. O’Meara TR, O’Meara MJ. 2021. DeORFanizing *Candida albicans* Genes using Coexpression. mSphere 6.

30. Ng H, Dean N. 2017. Dramatic Improvement of CRISPR/Cas9 Editing in *Candida albicans* by Increased Single Guide RNA Expression. mSphere 2.

31. Schiffer S, Rösch S, Marchfelder A. 2002. Assigning a function to a conserved group of proteins: the tRNA 3’-processing enzymes. EMBO J 21:2769–2777.

32. Gao Y, Zhao Y. 2014. Self-processing of ribozyme-flanked RNAs into guide RNAs in vitro and in vivo for CRISPR-mediated genome editing. J Integr Plant Biol 56:343–349.

33. Oud B, Guadalupe-Medina V, Nijkamp JF, de Ridder D, Pronk JT, van Maris AJA, Daran J-M. 2013. Genome duplication and mutations in *ACE2* cause multicellular, fast-sedimenting phenotypes in evolved *Saccharomyces cerevisiae*. Proc Natl Acad Sci U S A 110:E4223–31.

34. Kelly MT, MacCallum DM, Clancy SD, Odds FC, Brown AJP, Butler G. 2004. The *Candida albicans* Ca*ACE2* gene affects morphogenesis, adherence and virulence. Mol Microbiol 53:969–983.

35. Calderón-Noreña DM, González-Novo A, Orellana-Muñoz S, Gutiérrez-Escribano P, Arnáiz-Pita Y, Dueñas-Santero E, Suárez MB, Bougnoux M-E, Del Rey F, Sherlock G, d’Enfert C, Correa-Bordes J, de Aldana CRV. 2015. A single nucleotide polymorphism uncovers a novel function for the transcription factor Ace2 during *Candida albicans* hyphal development. PLoS Genet 11:e1005152.

36. Sreenivasan A, Kellogg D. 1999. The elm1 kinase functions in a mitotic signaling network in budding yeast. Mol Cell Biol 19:7983–7994.

37. Ito Y, Miyazaki T, Tanaka Y, Suematsu T, Nakayama H, Morita A, Hirayama T, Tashiro M, Takazono T, Saijo T, Shimamura S, Yamamoto K, Imamura Y, Izumikawa K, Yanagihara K, Kohno S, Mukae H. 2020. Roles of Elm1 in antifungal susceptibility and virulence in *Candida glabrata*. Sci Rep 10:9789.

38. King L, Butler G. 1998. Ace2p, a regulator of *CTS1* (chitinase) expression, affects pseudohyphal production in *Saccharomyces cerevisiae*. Curr Genet 34:183–191.

39. Schmidt M, Bowers B, Varma A, Roh D-H, Cabib E. 2002. In budding yeast, contraction of the actomyosin ring and formation of the primary septum at cytokinesis depend on each other. J Cell Sci 115:293–302.

40. Munro CA, Winter K, Buchan A, Henry K, Becker JM, Brown AJ, Bulawa CE, Gow NA. 2001. Chs1 of *Candida albicans* is an essential chitin synthase required for synthesis of the septum and for cell integrity. Mol Microbiol 39:1414–1426.

41. Saputo S, Chabrier-Rosello Y, Luca FC, Kumar A, Krysan DJ. 2012. The RAM network in pathogenic fungi. Eukaryot Cell 11:708–717.

42. Barrett-Bee K, Hamilton M. 1984. The detection and analysis of chitinase activity from the yeast form of Candida albicans. J Gen Microbiol 130:1857–1861.

43. Rybak JM, Muñoz JF, Barker KS, Parker JE, Esquivel BD, Berkow EL, Lockhart SR, Gade L, Palmer GE, White TC, Kelly SL, Cuomo CA, David Rogers P. 2020. Mutations in TAC1B: a Novel Genetic Determinant of Clinical Fluconazole Resistance in *Candida auris*. mBio.

44. Nelson B, Kurischko C, Horecka J, Mody M, Nair P, Pratt L, Zougman A, McBroom LDB, Hughes TR, Boone C, Luca FC. 2003. RAM: a conserved signaling network that regulates Ace2p transcriptional activity and polarized morphogenesis. Mol Biol Cell 14:3782–3803.

45. Du L-L, Novick P. 2002. Pag1p, a Novel Protein Associated with Protein Kinase Cbk1p, Is Required for Cell Morphogenesis and Proliferation inSaccharomyces cerevisiae. MBoC 13:503–514.

46. Dünkler A, Walther A, Specht CA, Wendland J. 2005. *Candida albicans CHT3* encodes the functional homolog of the Cts1 chitinase of Saccharomyces cerevisiae. Fungal Genet Biol 42:935–947.

47. Kuranda MJ, Robbins PW. 1991. Chitinase is required for cell separation during growth of *Saccharomyces cerevisiae*. J Biol Chem 266:19758–19767.

48. Sharma M, Chakrabarti A. 2020. On the Origin of *Candida auris:* Ancestor, Environmental Stresses, and Antiseptics. MBio 11.

49. Casadevall A, Kontoyiannis DP, Robert V. 2019. On the Emergence of *Candida auris*: Climate Change, Azoles, Swamps, and Birds. MBio 10.

50. Arora P, Singh P, Wang Y, Yadav A, Pawar K, Singh A, Padmavati G, Xu J, Chowdhary A. 2021. Environmental Isolation of *Candida auris* from the Coastal Wetlands of Andaman Islands, India. MBio 12.

51. Bartnicki-Garcia S. 1973. Fundamental aspects of hyphal morphogenesis, p. 245–267. In Microbe Differentiation 23rd Symposium of the Society of General Microbiology. Cambridge University Press.

52. Selvaggini S, Munro CA, Paschoud S, Sanglard D, Gow NAR. 2004. Independent regulation of chitin synthase and chitinase activity in *Candida albicans* and *Saccharomyces cerevisiae*. Microbiology 150:921–928.

53. Mondeel TDGA, Holland P, Nielsen J, Barberis M. 2019. ChIP-exo analysis highlights Fkh1 and Fkh2 transcription factors as hubs that integrate multi-scale networks in budding yeast. Nucleic Acids Res 47:7825–7841.

54. Greig JA, Sudbery IM, Richardson JP, Naglik JR, Wang Y, Sudbery PE. 2015. Cell cycle-independent phospho-regulation of Fkh2 during hyphal growth regulates *Candida albicans* pathogenesis. PLoS Pathog 11:e1004630.

55. Wakade RS, Ristow LC, Stamnes MA, Kumar A, Krysan DJ. 2020. The Ndr/LATS Kinase Cbk1 Regulates a Specific Subset of Ace2 Functions and Suppresses the Hypha-to-Yeast Transition in *Candida albicans*. MBio 11.

56. Koehler CM, Myers AM. 1997. Serine-threonine protein kinase activity of Elm1p, a regulator of morphologic differentiation in *Saccharomyces cerevisiae*. FEBS Lett 408:109–114.

57. Sutherland CM, Hawley SA, McCartney RR, Leech A, Stark MJR, Schmidt MC, Hardie DG. 2003. Elm1p is one of three upstream kinases for the *Saccharomyces cerevisiae* SNF1 complex. Curr Biol 13:1299–1305.

58. McCreath KJ, Specht CA, Robbins PW. 1995. Molecular cloning and characterization of chitinase genes from *Candida albicans*. Proc Natl Acad Sci U S A 92:2544–2548.

59. Shen J, Guo W, Köhler JR. 2005. CaNAT1, a heterologous dominant selectable marker for transformation of *Candida albicans* and other pathogenic *Candida* species. Infect Immun 73:1239–1242.

60. Walton FJ, Idnurm A, Heitman J. 2005. Novel gene functions required for melanization of the human pathogen *Cryptococcus neoformans*. Mol Microbiol 57:1381–1396.

61. Cowen LE, Singh SD, Köhler JR, Collins C, Zaas AK, Schell WA, Aziz H, Mylonakis E, Perfect JR, Whitesell L, Lindquist S. 2009. Harnessing Hsp90 function as a powerful, broadly effective therapeutic strategy for fungal infectious disease. Proc Natl Acad Sci U S A 106:2818–2823.

62. Hood EE, Gelvin SB, Melchers LS, Hoekema A. 1993. New Agrobacterium helper plasmids for gene transfer to plants. Transgenic Res 2:208–218.

63. Norrander J, Kempe T, Messing J. 1983. Construction of improved M13 vectors using oligodeoxynucleotide-directed mutagenesis. Gene.

64. Veri AO, Miao Z, Shapiro RS, Tebbji F, O’Meara TR, Kim SH, Colazo J, Tan K, Vyas VK, Whiteway M, Robbins N, Wong KH, Cowen LE. 2018. Tuning Hsf1 levels drives distinct fungal morphogenetic programs with depletion impairing Hsp90 function and overexpression expanding the target space. PLoS Genet 14:e1007270.

65. O’Meara TR, O’Meara MJ, Polvi EJ, Pourhaghighi MR, Liston SD, Lin Z-Y, Veri AO, Emili A, Gingras A-C, Cowen LE. 2019. Global proteomic analyses define an environmentally contingent Hsp90 interactome and reveal chaperone-dependent regulation of stress granule proteins and the R2TP complex in a fungal pathogen. PLoS Biol 17:e3000358.

66. Basenko EY, Pulman JA, Shanmugasundram A, Harb OS, Crouch K, Starns D, Warrenfeltz S, Aurrecoechea C, Stoeckert CJ Jr, Kissinger JC, Roos DS, Hertz-Fowler C. 2018. FungiDB: An Integrated Bioinformatic Resource for Fungi and Oomycetes. J Fungi (Basel) 4.

67. Esher SK, Granek JA, Alspaugh JA. 2015. Rapid mapping of insertional mutations to probe cell wall regulation in *Cryptococcus neoformans*. Fungal Genet Biol 82:9–21.

68. Afgan E, Baker D, Batut B, van den Beek M, Bouvier D, Cech M, Chilton J, Clements D, Coraor N, Grüning BA, Guerler A, Hillman-Jackson J, Hiltemann S, Jalili V, Rasche H, Soranzo N, Goecks J, Taylor J, Nekrutenko A, Blankenberg D. 2018. The Galaxy platform for accessible, reproducible and collaborative biomedical analyses: 2018 update. Nucleic Acids Res 46:W537–W544.

69. Martin M. 2011. Cutadapt removes adapter sequences from high-throughput sequencing reads. EMBnet.journal 17:10–12.

70. Li H, Durbin R. 2009. Fast and accurate short read alignment with Burrows-Wheeler transform. Bioinformatics.

71. Madeira F, Park YM, Lee J, Buso N, Gur T, Madhusoodanan N, Basutkar P, Tivey ARN, Potter SC, Finn RD, Lopez R. 2019. The EMBL-EBI search and sequence analysis tools APIs in 2019. Nucleic Acids Res 47:W636–W641.

72. Schindelin J, Arganda-Carreras I, Frise E, Kaynig V, Longair M, Pietzsch T, Preibisch S, Rueden C, Saalfeld S, Schmid B, Tinevez J-Y, White DJ, Hartenstein V, Eliceiri K, Tomancak P, Cardona A. 2012. Fiji: an open-source platform for biological-image analysis. Nat Methods 9:676–682.

73. Lee DW, Hong CP, Kang HA. 2019. An effective and rapid method for RNA preparation from non-conventional yeast species. Anal Biochem 586:113408.

74. Aranda PS, LaJoie DM, Jorcyk CL. 2012. Bleach gel: a simple agarose gel for analyzing RNA quality. Electrophoresis 33:366–369.

